# DNA metabarcoding from sample fixative as a quick and voucher preserving biodiversity assessment method

**DOI:** 10.1101/287276

**Authors:** Vera Marie Alida Zizka, Florian Leese, Bianca Peinert, Matthias Felix Geiger

**Affiliations:** Aquatic Ecosystem Research, Faculty of Biology, University of Duisburg-Essen, Universitätsstraße 5, 45141 Essen, Germany; Centre for Water and Environmental Research (ZWU) Essen, University of Duisburg-Essen, Universitätsstraße 2, 45141 Essen, Germany; Zoologisches Forschungsmuseum Alexander Koenig, Leibniz Institute for Animal Biodiversity, Adenauerallee 160, 53113 Bonn, Germany

**Keywords:** non-destructive, environmental DNA, macroinvertebrates, metabarcoding

## Abstract

Metabarcoding is a powerful tool for biodiversity assessment and has become increasingly popular in recent years. Although its reliability and applicability have been proven in numerous scientific studies, metabarcoding still suffers from some drawbacks. One is the usually mandatory destruction of specimens before DNA extraction, which is problematic because it does not allow a later taxonomic evaluation of the results. Additionally, metabarcoding often implements a time-consuming step, where specimens need to be separated from substrate or sorted in different size classes. A non-destructive protocol, excluding any sorting step, where the extraction of DNA is conducted from a samples fixative (ethanol) could serve as an alternative. We test an innovative protocol, where the sample preserving ethanol is filtered and DNA extracted from the filter for subsequent DNA metabarcoding. We first tested the general functionality of this approach on 15 mock communities comprising one individual of eight different macroinvertebrate taxa each and tried to increase DNA yield through different treatments (ultrasonic irradiation, shaking, freezing). Application of the method was successful for most of the samples and taxa, but showed weaknesses in detecting mollusc taxa. In a second step, the community composition detected in DNA from ethanol was compared to conventional bulk sample metabarcoding of complex environmental samples. We found that especially taxa with pronounced exoskeleton or shells (Coleoptera, Isopoda) and small taxa (Trombidiformes) were underrepresented in ethanol samples regarding taxa diversity and read numbers. However, read numbers of Diptera (mainly chironomids) and Haplotaxida were higher in ethanol derived DNA samples, which might indicate the detection of stomach content, which would be an additional advantage of this approach. Concerning EPT (Ephemeroptera, Plecoptera, Trichoptera) taxa which are decisive for the determination of ecological statuses, both methods had 46 OTUs in common with 4 unique to the ethanol samples and 10 to the bulk samples. Results indicate that fixative-based metabarcoding is a non-destructive, time-saving alternative for biodiversity assessments focussing on taxa used for ecological status determination. For a comprehensive picture on total biodiversity, the method might however not be sufficient and conventional bulk sample metabarcoding should be applied.

## 1. Introduction

DNA metabarcoding was developed to improve the efficiency, reliability and resolution of biodiversity assessments. The technique is based on parallel sequencing of standardised DNA barcode markers from several to thousands of organisms in one sample. Compared to traditional morphology-based assessments, metabarcoding has the clear advantage of higher taxonomic resolution, better comparability and often also speed (Haase et al. 2010, Baird and Hajibabaei 2012, Hajibabaei et al. 2012, Taberlet et al. 2012, Elbrecht et al. 2017). Even though metabarcoding has been successfully applied for biodiversity assessments (e. g., Shokralla et al. 2012, Yu et al. 2012, Leray and Knowlton 2014, Elbrecht et al. 2017), the method still lacks consistent protocols (Baird and Hajibabaei 2012, Taberlet et al. 2012, Leese et al. 2016, Leese et al. 2018). Furthermore, one particular drawback of metabarcoding when working with complex environmental samples (e. g. invertebrates from the streambed, so-called bulk metabarcoding) is that organisms have to be separated manually from the by-catch (sediment and other inorganic matter) prior to DNA extraction. This is often time-consuming and thus limits the gain in speed substantially (Elbrecht et al. 2017a). Furthermore, as specimens are usually homogenised and destroyed for DNA extraction they consequently cannot be used for further morphological examination or validation (Zimmermann et al. 2008, Leese et al. 2016). The non-destructive isolation of DNA from the bulk samples’ fixative (most often ethanol) has been put forward as a promising alternative (Hajibabaei et al. 2012). However, studies following this idea still included sample sorting and/or the usage of lysis buffer plus proteinase to soften body structures of specimens for an increased DNA release. This can lead to the complete destruction of smaller specimens or of those lacking a strong internal or external skeleton, again prohibiting subsequent morphological identification (Shokralla et al. 2010, Hajibabaei et al. 2012, Taberlet et al. 2012).

In the present study we advanced the state of the art of metabarcoding from preservation ethanol as proposed by Hajibabaei et al. (2012) by extracting DNA directly from the ethanol with a novel filter-based approach. This approach does not include any preceding tissue digestion to maximize morphological integrity of specimens. We tested the approach on pre-sorted small mock communities of invertebrates and on real environmental samples that included substrates (e.g. sand, stones, litter etc.), potentially also acting as PCR inhibitors. First, the general feasibility of ethanol filtration and subsequent extraction of DNA was evaluated using 15 different mock communities with known taxonomic composition to address 1) if DNA release is high enough to obtain sufficient DNA to correctly detect the mock community composition, 2) if different treatments of samples (ultrasonic irradiation, shaking, freezing) could increase DNA release and thereby detection success, and 3) if final read abundance is correlated with biomass or size of specimens. We hypothesized that larger specimens are less overrepresented in read numbers based on DNA from ethanol than including homogenization of specimens prior to metabarcoding (Elbrecht and Leese 2015) given that the surface area to volume ratio decreases with increased specimen size.

In the second part of the study, we analysed real environmental samples by the same metabarcoding pipeline: we used the ethanol phase of six aquatic multi-habitat environmental samples and compared the results to a conventional tissue-homogenisation-based metabarcoding protocol (e. g., Elbrecht et al. 2017a). Specifically, we evaluated if it is possible to detect similar biodiversity and community compositions through the extraction of DNA from the fixative and subsequent metabarcoding as with the tissue-homogenisation-based metabarcoding protocol.

Addressing these points in a controlled comparative framework is essential in order to validate if using the fixative for metabarcoding can become part of an accelerated bioassessment protocol. The chosen experimental design enables the validation of fixative-based metabarcoding which can be used for further method improvements.

## 2. Material and Methods

### 2.1 Testing filtration of ethanol and isolation of DNA from filters

#### 2.1.1 Assembly of mock communities

For part one of this study, fifteen mock communities were assorted in November 2016 directly *in situ* at the Deilbach in Velbert, North Rhine Westphalia, Germany. Each community contained one single specimen of the eight morphotaxa *Ancylus*, *Ecdyonurus*, *Ephemera, Gammarus*, Gastropoda (*Potamopyrgus* or *Radix*), *Hydropsyche*, Leptophlebiidae and *Polycentropus*, respectively. Size relations between specimens were tried to be kept similar for three mock communities each (e. g., A always S[mall]+L[large]+M[edium]+L+M+L+S+M). Three communities of the same size composition were then treated as replicates (e. g. A1-A3) for further analyses and 5 communities of different size relations were assembled (A-E; Fig. 1, Tab. S1), resulting in 15 physical mock samples. Specimens were collected and directly transferred to a falcon tube containing 50 ml of 96 % denatured technical ethanol (410 PharmEur., ethanol phase I). Samples were transported to the laboratory of the University of Duisburg-Essen and stored for 12 hours at −20 °C. After 12 hours, the ethanol (phase I) was poured over a sieve (mesh size 0.5 mm) to retain animals and substrate and was then stored at −20 °C until further processing (protocol 0, see section 2.1.3 Laboratory protocols filtration and DNA extraction). Falcon tubes with the respective communities were filled up with 50 ml of denatured technical 96 % ethanol (410 PharmEur. ethanol phase II) and stored at −20 °C until further processing.

**Figure 1:**
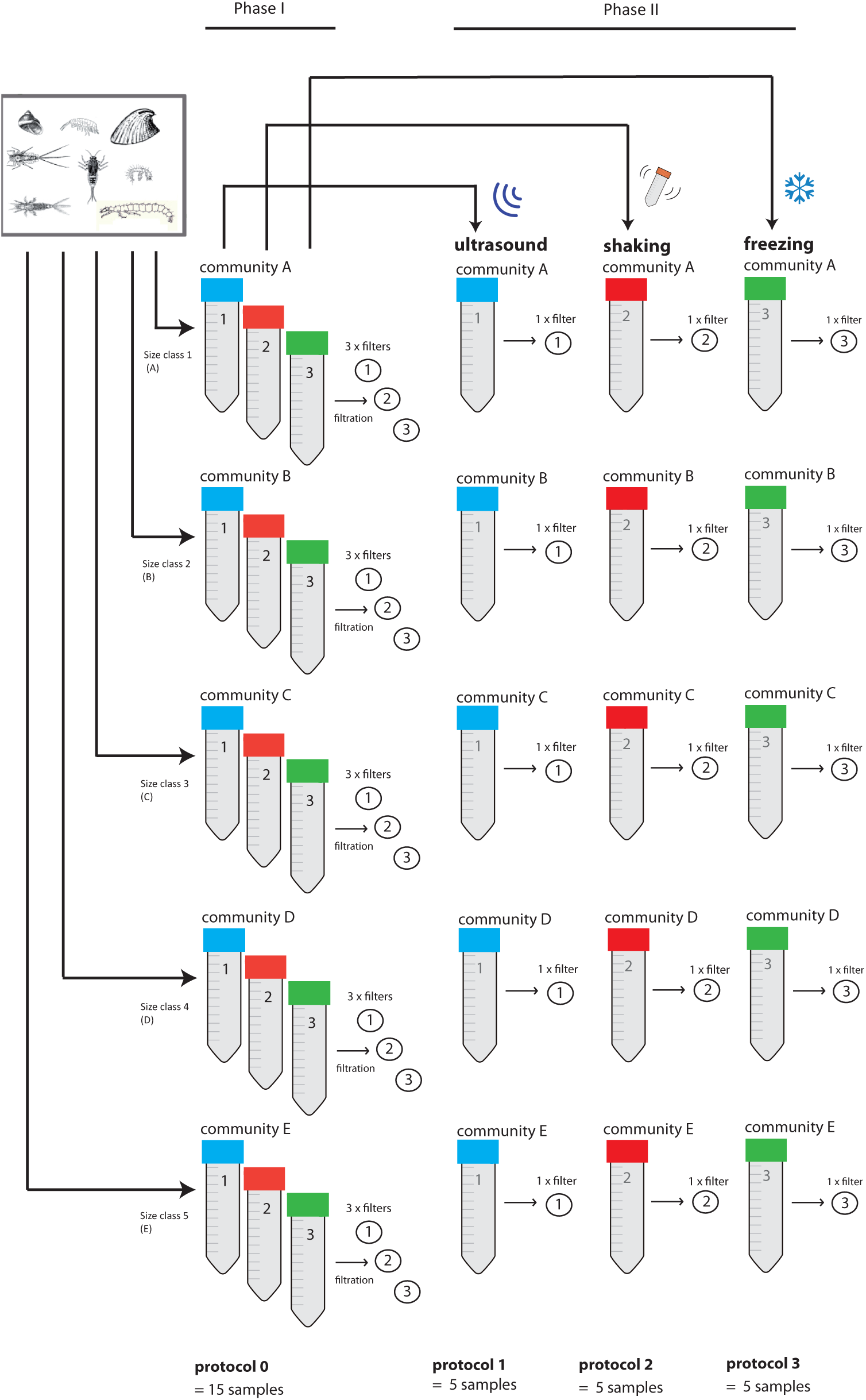
In total, 15 different mock communities composed of eight specimens (one individual each from the following taxa: *Ancylus*, *Ecdyonurus*, *Ephemera, Gammarus*, Gastropoda (*Potamopyrgus* or *Radix*), *Hydropsyche*, Leptophlebiidae and *Polycentropus*) were assembled. Individuals of very similar sizes per taxon were compiled for each of the three communities to allow for comparisons of extraction efficiencies among these three replicates (e. g., A1-A3). The ethanol in which specimens were initially preserved after collection was filtrated for all 15 communities (protocol 0 à Phase I). The second ethanol phase was then filtrated after one of the three treatments: ultrasonic irradiation, shaking or freezing (protocol 1-3).

**Table 1:** Proportion of total reads [%] assigned to the eight taxa of the mock communities. Protocols: 0 = first ethanol phase in which specimens were preserved after collection; 1 = second ethanol phase and treatment with ultrasonic irradiation; 2 = second ethanol phase and shaking; 3 = second ethanol phase and freezing. Question marks show cases where taxonomic assignments through single-specimen barcoding could not be validated in the metabarcoding dataset.

| Community | Protocol | *Ancyclus* | *Ecdyonurus* | *Ephemera* | *Gammarus* | Gastropoda | *Hydropsyche* | *Leptophlebiidae* | *Polycentropus* | Total [%] |
| --- | --- | --- | --- | --- | --- | --- | --- | --- | --- | --- |
| A1 | 0 | 0 | 0.24 | 0.724 | 12.156 | 0.452 | 17.251 | 0.578 | 0.001 | 31.402 |
| A2 | 0 | 0.011 | 0.037 | 1.49 | 0.124 | 0.017 | 26.782 | 0 | 0.055 | 28.516 |
| A3 | 0 | 0.001 | 0.156 | 0.108 | 0.329 | 0.091 | 28.638 | 0.36 | 0.043 | 29.726 |
| B1 | 0 | 0.001 | 0.182 | 0.375 | 1.452 | 0 | 25.343 | ? | 0.044 | 27.397 |
| B2 | 0 | 0.003 | 0.114 | 0.309 | 1.778 | 0.046 | 32.929 | 3.998 | 0.185 | 39.362 |
| B3 | 0 | 0 | 1.304 | 0.014 | 0.149 | 0 | 38.323 | 0.023 | 0.669 | 40.482 |
| C1 | 0 | 0 | 8.311 | 8.737 | 0.762 | ? | 1.084 | 0.269 | 0.156 | 19.319 |
| C2 | 0 | 0.009 | 0.019 | 1.329 | 1.801 | 0.328 | 0 | 0 | 0 | 3.486 |
| C3 | 0 | 0 | 2.201 | 0 | 0.163 | 1.212 | 6.205 | 0.734 | 0.051 | 10.386 |
| D1 | 0 | 0 | 0.006 | 0.298 | 8.931 | 0 | 0.768 | 0.367 | 0.216 | 10.586 |
| D2 | 0 | 0.002 | 0.054 | 80.586 | 1.109 | 0.174 | 0.692 | 0.696 | 0.022 | 83.272 |
| D3 | 0 | 0.011 | 0.093 | 0.255 | 0.358 | 0.064 | 77.913 | 0.088 | 0.625 | 79.407 |
| E1 | 0 | 0.013 | 0.869 | 5.198 | 2.789 | 0 | 0.27 | 0 | 0.126 | 9.265 |
| E2 | 0 | 0.007 | 1.312 | 33.267 | 3.994 | 0.061 | 9.093 | 0.269 | 0.956 | 48.959 |
| E3 | 0 | 0 | 3.889 | 0.333 | 0.115 | 0.148 | 1.301 | 3.932 | 35.046 | 44.764 |
| A1 | 1 | 0 | 0 | 0 | 0 | 0 | 0 | 0 | 0 | 0 |
| B1 | 1 | 0 | 0 | 0 | 0 | 0 | 0 | 0 | 0 | 0 |
| C1 | 1 | 0 | 0 | 0 | 0 | 0 | 0 | 0 | 0 | 0 |
| D1 | 1 | 0 | 0 | 0 | 0 | 0 | 0 | 0 | 0 | 0 |
| E1 | 1 | 0 | 0 | 0 | 0 | 0 | 0 | 0 | 0 | 0 |
| A2 | 2 | 0 | 49.238 | 0.838 | 14.968 | 0.002 | 14.968 | 7.248 | 0 | 87.262 |
| B2 | 2 | 0.003 | 0.078 | 0.703 | 0.751 | 0.019 | 10.579 | 37.129 | 0 | 49.262 |
| C2 | 2 | 0.006 | 0.809 | 38.681 | 1.69 | 0.026 | 0.034 | 16.862 | 0.02 | 58.298 |
| D2 | 2 | 0.003 | 0.034 | 47.076 | 0.398 | 0.076 | 1.015 | 39.326 | 0 | 87.928 |
| E2 | 2 | 0 | 0 | 0 | 0 | 0 | 0 | 0 | 0 | 0 |
| A3 | 3 | 0 | 1.154 | 53.825 | 1.508 | 0.008 | 10.999 | 24.392 | 0.188 | 93.074 |
| B3 | 3 | 0 | 45.056 | 12.484 | 4.494 | 0 | 9.52 | 3.904 | 16.346 | 91.804 |
| C3 | 3 | 0 | 0.985 | 78.12 | 0.022 | 0.001 | 0.436 | 17.587 | 0.004 | 97.155 |
| D3 | 3 | 0 | 2.162 | 0.06 | 4.513 | 0.004 | 18.509 | 63.461 | 0.066 | 88.775 |
| E3 | 3 | 0 | 0.092 | 31.456 | 8.848 | 0.001 | 0.092 | 14.741 | 42.551 | 97.781 |

#### 2.1.2 Treatments

After the first ethanol change, the mock communities underwent three different treatments (Figure 1 protocol 1-3) to potentially increase DNA release into the ethanol. From the five differently composed (in terms of specimen size) mock communities, each one of three replicates was used per treatment (protocol 1–3; see Fig. 1, Tab. S1):

**Protocol (1)** ultrasonic irradiation: indirect ultrasonic irradiation for 15 minutes at 35 kHz (Bandalin SONOREX Super rk 510 Hz).
**Protocol (2)** mechanical shaking: shaking for 2 minutes at 2000 rpm at room temperature using the Eppendorf ThermoMixer C.
**Protocol (3)** freezing: samples were kept for 10 minutes in liquid nitrogen (−195.79°C) until ethanol was frozen. Afterwards, samples were left for 15 minutes at room temperature for thawing. The procedure was repeated three times.

After each treatment the ethanol (phase II) was filtered through a sieve (mesh size 0.5 mm) to retain and was stored at −20 °C until further processing (protocols 1-3, see section 2.1.3 Laboratory protocols filtration and DNA extraction).

#### 2.1.3 Laboratory protocols

Ethanol was filtered through nitrocellulose filters with pore size 0.45 μm (Nalgene Analytical Test Filter Funnel CN, Thermo Scientific) using a vacuum pump (TC-501v, Sparmax). Filtration, as well as all following laboratory processes (except treatments) were conducted in a sterile lab, set up for the treatment of eDNA (environmental DNA) samples and complete body protection (overalls with hood, mouth protection, gloves) was worn at all laboratory steps. Filters were dried overnight in petri dishes. Dry filters were ripped into small pieces using sterile plastic tweezers and transferred to 600 µL TNES buffer for subsequent DNA extraction. DNA extraction was carried out using a modified salt extraction protocol (Sunnucks and Hales 1996, adjusted in Weiss and Leese 2016). Extraction success was checked on a Fragment Analyzer (Advanced Analytical) and 1 μL of each sample was used for amplicon library preparation in a two-step PCR approach targeting a 421 bp long fragment of the official animal DNA barcode region (5’ end of the mitochondrial cytochrome C oxidase subunit (CO1) gene). In the first step the fragment was amplified using untailed BF2/BR2 primers (Elbrecht and Leese 2017). PCR reactions consisted of 1 x PCR buffer (including 2.5 mM Mg^2+^), 0.2 mM dNTPs, 0.5 µM of each primer, 0.025 U/μL of HotMaster Taq (5 Prime, Gaithersburg, MD, USA), 1 μl DNA template filled up with HPLC H_2_O to a total volume of 50 μl. The following PCR program was used: 94 °C for 180 s, 25 cycles of 94 °C for 30 s, 50 °C for 30 s and 65 °C for 150 s followed by a final elongation of 65 °C for 5 min at a Thermocycler (Biometra TAdvanced Thermocycler). For the second PCR step, 1 μl of each PCR product was used in order to add labels using uniquely indexed BF2/BR2 fusion primers for pooling the samples into one amplicon library for subsequent sequencing. PCR conditions in the second step were identical as in the previous step but only 15 cycles were used.

A left sided size selection of all samples was performed using magnetic beads (SPRIselect BECKMAN COULTER), to clean samples from short fragments (leftover primers and primer dimers). A ratio of 0.76x was applied and amplicon concentration was measured on a Fragment Analyzer (Advanced Analytical). All samples were equimolarly pooled and paired-end sequencing was carried out by GATC Biotech AG (Constance, Germany) using one run on the Illumina MiSeq platform with a 250 bp paired-end v2 kit.

#### 2.1.4 Sanger sequencing of specimens

In addition to DNA extraction from filters and subsequent sequencing from filters, full length DNA barcodes of all 120 individuals of the 15 mock communities were obtained through the German Barcode of Life (GBOL) pipeline at ZFMK in order to allow direct comparisons of OTUs to the specimens. For single specimen DNA barcoding, genomic DNA was extracted from sub-samples using a BioSprint96 magnetic bead extractor (Qiagen, Hilden, Germany). PCRs were carried out in 20 μl reaction volumes including 2 μl undiluted DNA template, 0.8 μl of each primer (10 pmol/μl; LCO1490-JJ: 5’-CHACWAAYCATAAAGATATYGG-3’ and HCO2198-JJ: 5’-AWACTTCVGGRTGVCCAAARAATCA-3’, Astrin and Stüben 2008), 2 μl ‘Q-Solution’ and 10 μl ‘Multiplex PCR Master Mix’ (Qiagen, Hilden, Germany). Thermal cycling was performed on GeneAmp PCR System 2700 machines (Applied Biosystems, Foster City, CA, USA) as follows: hot start Taq activation: 15 min at 95°C; first cycle set (15 repeats): 35 s denaturation at 94°C, 90 s annealing at 55°C (−0.2°C/cycle, ‘touch down’) and 90 s extension at 72°C. Second cycle set (25 repeats): 35 s denaturation at 94°C, 90 s annealing at 50°C and 90 s extension at 72°C; final elongation 10 min at 72°C. Purification of PCR products and sequencing in both directions was conducted at BGI (Hongkong, China) using the amplification primers.

#### 2.1.5 Data Analysis

For the metabarcoding approach, sequences labelled with fusion primers and indices were assigned to their original sample as implemented in JAMP v0.23 (https://github.com/VascoElbrecht/JAMP). Subsequent data processing was conducted for all samples in JAMP v0.23 using standard settings: paired-end reads were first merged (module U_merge_PE) and reverse complements were built where needed (U_revcomp) with usearch v10.0.240 (Edgar 2010). Cutadapt (Martin 2017) was used to remove primers and to discard sequences of unexpected length so that only reads with a length of 231 – 251 bp were used for further analyses (Minmax()). The module U_max_ee was used to discard all reads with an expected error > 0.5. Sequences were dereplicated, singletons were removed and sequences with ≥ 97 % similarity were clustered into OTUs using Uparse (U_cluster_otus). OTUs with a minimal read abundance of 0.01 % in at least one sample were retained for further analyses while other OTUs were discarded. The used script for data analysis can be found in supplement S1. OTU sequences were compared with the database BOLD using the BOLD ID engine via the module BOLD_webhack in JAMP and taxonomies assigned following rules outlined in Elbrecht et al. 2017: for assignment to species level, a hit with 98% similarity was required; 95% similarity was required for assignment to genus level, 90% for family level, and 85% for order level. OTU sequences were also compared to the sequences generated by single specimen DNA barcoding to assess the number of regained taxa and the general success of fixative-based metabarcoding.

### 2.2 Comparison of ethanol samples with environmental bulk samples

#### 2.2.1 Sampling

For part two of this study, we compared the performance of fixative-based metabarcoding to tissue-homogenisation-based metabarcoding of six real environmental samples. Therefore, six sites along the river Sieg were visited in spring 2017 and a multi-habitat sampling was conducted according to the Water Framework Directive (WFD) guidelines (Meier et al. 2006), i.e. by taking 20 subsamples (kick-sampling) at each sampling site covering all different microhabitats. Complete samples (containing benthic macroinvertebrates and substrate) were transferred to 96% denatured technical ethanol. For each sampling site, two bottles (1 litre) were needed to capture the whole bulk sample. Immediately upon arrival at the laboratory (ca. 3-6 hrs after sampling), the ethanol of all samples was poured off through a sieve (mesh size: 0.5 mm) and stored at −20 °C. New 96% denatured technical ethanol was added to the samples, which were then also stored at −20 °C until further processing.

#### 2.2.2 Processing of ethanol samples

The following laboratory steps were carried out for both bottles per sample site separately. Ethanol filtration of the first phase used for fixation in the field was performed as outlined above for the mock communities and filters were dried overnight in petri dishes. Dry filters were ripped into small pieces using sterile plastic tweezers and transferred to 600 µL TNES buffer for subsequent DNA extraction. Both PCR steps were conducted with Illustra^TM^ PuRe Taq Ready-To-go^TM^ PCR beads. Per sample, 0.5 μM BF2 and BR2 primer were added to an Illustra bead and filled up to a total volume of 25 µL with HPLC H_2_O. The following PCR program was used for the first PCR step: 95°C for 180 s, 25 cycles of 95°C for 30 s, 48°C for 30 s and 72°C for 60 s followed by a final elongation of 72°C for 5 min at a Thermocycler (Biometra TAdvanced Thermocycler). The second PCR step was conducted with similar conditions but with an annealing temperature of 50 °C and 15 cycles. All following steps were carried out as described in chapter 2.1.3, Laboratory protocols.

#### 2.2.3 Processing of environmental bulk samples

All macroinvertebrates in each sample were separated from the substrate, counted and categorised according to their size into two classes using standardized reference areas of 5 mm x 2 mm. Individuals fitting into this area were assigned to size class S, individuals exceeding this area were assigned to size class L. All specimens of the respective size class were homogenised to fine powder with an IKA Ultra Turrax Tube Disperser (full speed for 30 minutes) and DNA was extracted following a modified salt extraction protocol (Sunnucks and Hales 1996, adjusted as in Weiss and Leese 2016). After DNA extraction, the two size categories of the same sample were pooled in proportion to individual numbers (e. g., if 90 % of individuals were assigned to size class S and 10 % to size class L, for a 20 μl dilution 18 μl of the extract of size class S was pooled with 2 μl of size class L, see Elbrecht et al. 2017a for the detailed description). DNA concentration was quantified using a Qubit 2.0 Fluorometer (Life technologies, dsDNA BR Assay Kit) and DNA diluted to 25 ng/μl. A two-step PCR (see section 2.1.3) was conducted for all samples with one PCR replicate each. The PCR products were quantified using a Fragment Analyzer (Advanced Analytical) and all samples were pooled equimolarly. Samples were sent to GATC Biotech (Constance, Germany) and sequenced on an Illumina MiSeq platform with a 250 bp paired-end v2 kit.

#### 2.2.4 Data Analysis

Data were analysed using the program JAMP v0.23 (https://github.com/VascoElbrecht/JAMP) as described in section 2.1.5. To compare the fixative-based and bulk-sample metabarcoding, the obtained community compositions was visualized using Principal Coordinates analyses (PCoA) implemented PAST v. 3.17 (Hammer et al. 2001) based on Bray–Curtis (read abundance included) and Sørensen (presence-absence) similarities. Further data were visualised using the program R (R Development Core Team 2008) with the package asbio (Aho 2014) and ggplot2 (Wickham 2009).

## 3. Results

### 2.1 Testing filtration of ethanol and isolation of DNA from filters on mock communities

All raw data will be available on Short Read Archive, Submission number: XX. Extraction of DNA from ethanol was successful for all protocols except protocol 1 (ultrasonic irradiation) where no DNA product was visible on an agarose gel. However, samples were sent for sequencing as an additional validation that no product was present. A total of 10,659,925 raw reads were obtained from the Illumina MiSeq sequencing run. After demultiplexing, samples obtained from filtering the first ethanol phase (protocol 0, n=15) comprised on average 325,808 (± 170,687) reads. Samples treated with protocol 2 (shaking, n=5) and protocol 3 (freezing, n=5) contained on average 347,637 (± 236,007) and 456,342 (± 279,465) reads. Samples treated with protocol 1 (ultrasonic irradiation, n=5) contained on average only 49 (± 28.86) reads and were therefore excluded from further analyses (Tab. S2). In total 4583 OTUs were detected in this dataset with 950 OTUs assigned to macroinvertebrate taxa. The majority of OTUs were assigned to bacteria and algae.

Comparison of fixative-based metabarcoding with generated sequences through single-specimen barcoding revealed the recovery of 6.9 (± 0.99, n=15) detected morphotaxa for protocol 0 (Tab. S3). On average 33.76 % (± 23.7) of the total reads of each replicate were assigned to OTUs matching the Sanger sequencing-based eight target morphotaxa. The taxon that went undetected most often was *Ancylus fluviatilis*, which could not be detected in five of the 15 mock communities using protocol 0. Sequences of *Ancylus fluviatilis* were also rare in the other ten mock communities, where on average only 0.006 % of the total reads per community were assigned to this taxon (Tab. 1, Tab. S3). Single specimen barcoding revealed that the gastropods collected in the field belonged to different genera or species (Tab. S3). In four of the 15 mock communities those taxa could not be detected through fixative-based metabarcoding. For community B1 the specimen was identified as *Physa* sp. through single specimen barcoding. The sequence showed no similarity to any OTU sequence assigned to a mollusc taxa. This was similar for the species *Galba truncatula* which was assigned to the morphotaxon Gastropoda in community C1 through single specimen barcoding. For community B3, D1 and E1 the gastropod sequences could only be assigned to family level (Succineidae, terrestrial pulmonate gastropod molluscs) through single specimen barcoding, but could not be detected through the metabarcoding approach. In the other communities, taxa assigned to the morphotaxon Gastropoda (*Potamopyrgus antipodarum*, *Radix labiata*, *Physa acuta, Succinea putris*) contributed on average 0.26 % to the total read number. Beside the two mollusc taxa, which could not be found in several mock communities through fixative-based metabarcoding, only a few arthropods remained undetected in certain mock communities (A2: *Habroleptoides confusa*, C2: *Polycentropus flavomaculatus* and *Hydropsyche saxonica*, C3: *Ephemera danica*, E1: *Ecdyonurus torrentis*). In community B1 the mayfly *Baetis vernus* was identified through single specimen barcoding, but no OTU was assigned to this species through metabarcoding. No other Baetidae was highly abundant in this community (Tab. S5). Further 19.12 % (± 9.53) of the reads were on average assigned to other macroinvertebrate taxa, containing 56 dipteran OTUs. Chironomidae, Simulidae and Tipulidae were the dominant taxa represented by these OTUs (Tab. S6).

For protocol number 2 (shaking) on average 7 (± 0.82) of the morphotaxa were detected through the fixative-based metabarcoding. After bioinformatic analyses, sample E2 showed only 358 reads and was therefore excluded from further analyses. On average 70.65 % (± 19.9) of the total reads were assigned to the eight target taxa (Tab. S5). Additional aquatic macroinvertebrate taxa were detected with on average 9.35 % (± 7.2) of the total reads (Tab. S6). Only *Ancylus fluviatilis* (A2) and *Polycentropus flavomaculatus* (A2, B2, D2) could not be detected in some communities (Tab. S3).

For protocol number 3 (freezing) on average 6.8 (± 0.45) morphotaxa were detected. With on average 93.7 % (± 3.5) of the total read number being assigned to the eight target taxa (Tab. S3), this protocol had the lowest number of non-target sequence reads. Only 2.42 (± 1.36) % of the total reads were assigned to other macroinvertebrate taxa not contained in the mock communities (Tab. S6). However, in four of the five samples treated with protocol 3, *Ancylus fluviatilis* could not be detected and in sample B3 Succineidae sp. was also not recovered.

Only for protocol 0 a weak correlation between biomass and read numbers existed (Spearman’s rho 0.211, *p*=0.02). The strong scattering of the underlying data points and the fact that no significant correlation between biomass and read numbers was evident in the two other treatments indicate that biomass of individuals has an inferior effect on the final read number. However, the present results show that individuals release different amounts of DNA, which biases in turn the generated read numbers. For fixative-based metabarcoding, these biases seem to be more dependent on morphology of the specimens or on the random dispersal of cells or even body structures on the filter. An additional filtering with a finer mesh size than applied in the present study (0.5 mm) of the ethanol before processing could help circumventing the problem of complete cells on the filter, as recently also proposed by a comparison of different filtration techniques for eDNA from water samples (Majaneva et al. 2018).

### 2.2 Comparison of environmental bulk samples with ethanol samples

Altogether 8,879,690 reads were obtained from the ethanol samples and 8,545,029 reads from the bulk samples (including samples of another river system (Emscher), which were pooled for sequencing with the Sieg bulk samples). After demultiplexing and quality filtering of the derived ethanol samples 5,047,373 (average 210,307 ± 144,450) reads were retained. DNA extractions from one bottle sampled at sites S4 and S6 yielded less than 400 sequences and were therefore excluded from further analyses (Tab. S7). Sequencing of the second bottle from both sample sites showed a mean read number of 90,146 and 443,841 raw sequences, respectively. The bulk sample based DNA extraction resulted in 2,334,707 sequences (average: 194,558 ± 43,135) with no samples containing less than 30,000 reads. For further analyses, only OTUs present in both extraction and PCR replicates per sample site were kept in the dataset, and read numbers were added up to one OTU table. After this filtering step, ethanol samples included on average 836,027 (± 494,981) reads while bulk samples included on average 388,867 (± 74,318) reads (Fig. 2, Tab. S8).

**Figure 2:**
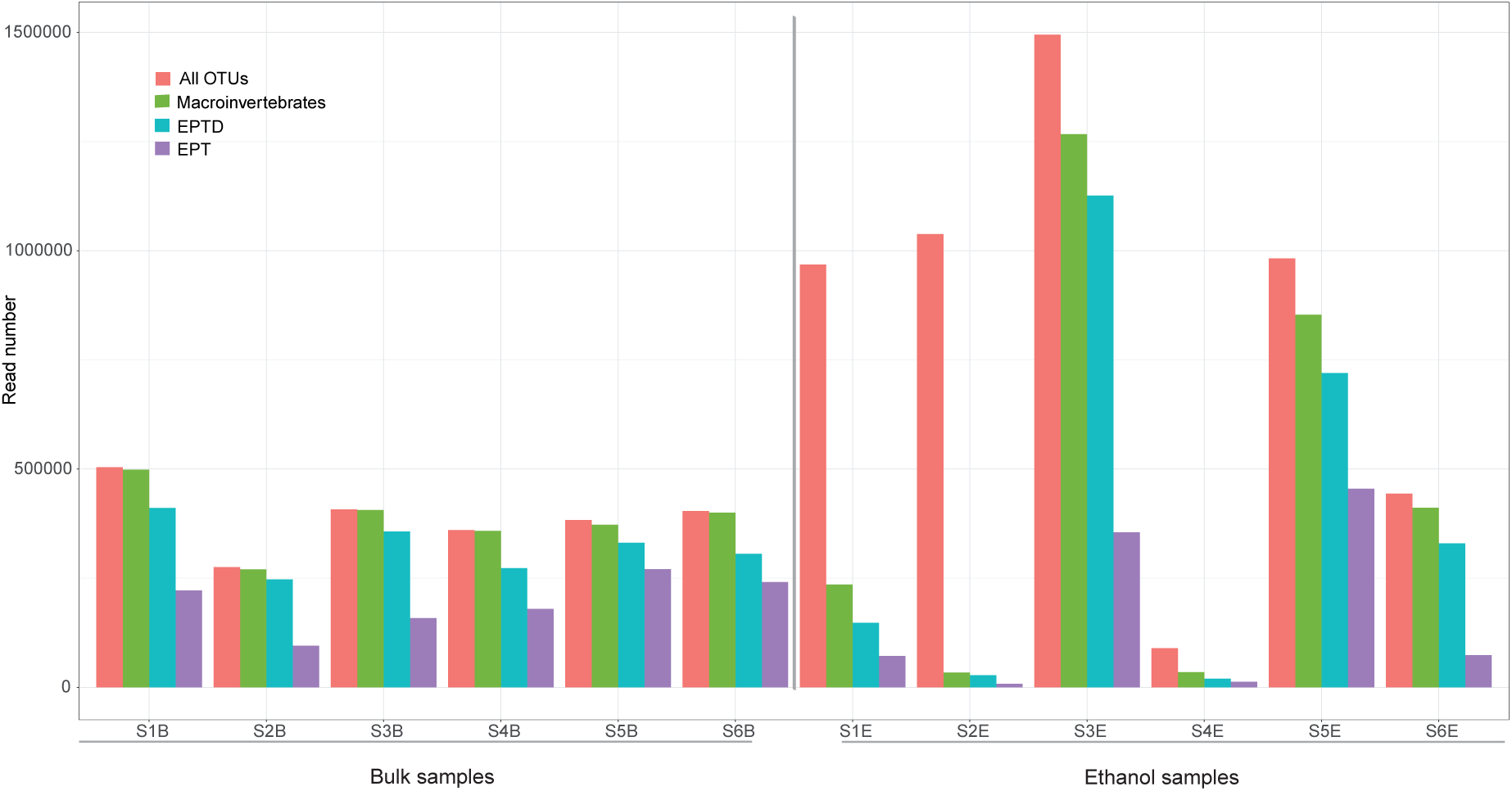
Read numbers per sample site (S1B-S6B = bulk samples, S1E-S6E = ethanol samples). Data include *all OTUs* with a threshold of 0.01 % of the total reads in at least one sample. No further filtering threshold was applied. *Macroinvertebrates* include all OTUs assigned to the taxa Arthropoda, Annelida, Mollusca and Platyhelminthes with at least 85 % similarity to a sequence deposited in BOLD. The category *EPTD* includes all OTUs assigned to the taxa Ephemeroptera, Plecoptera, Trichoptera and Diptera with at least 85 % similarity compared with the BOLD database. The category *EPT* includes the same taxa except Diptera.

Comparing OTUs among the two methods revealed a consensus of 357 OTUs, while 640 were unique to the ethanol and 104 to the bulk samples, respectively. Out of the OTUs uniquely found in ethanol samples the main proportion was assigned to algae (Ochrophyta) (Tab. 2, Tab. S8). OTUs uniquely found in bulk sample metabarcoding included several beetles (Coleoptera), caddisflies (Trichoptera), as well as few other aquatic macroinvertebrate taxa (Fig. 3, Tab. S8).

**Table 2:** Overview of the different datasets used for the analyses. Complete dataset includes (1) all OTUs above 0.01 % of the total reads in at least one sample and (2) OTUs with at least 85 % similarity to a sequence deposited in BOLD. Macroinvertebrates include all OTUs assigned to any taxa of Arthropoda, Annelida, Mollusca or Platyhelminthes with at least 85 % similarity to a sequence deposited in BOLD. The subsample includes only a random 230,000 reads of macroinvertebrates per sample, where S2 and S4 needed to be excluded due to low read numbers. EPTD taxa include all OTUs assigned to Ephemeroptera, Plecoptera, Trichoptera and Diptera with at least 85 % similarity to a sequence deposited in BOLD. Same holds true for the EPT taxa with Diptera excluded.

| Dataset | Included samples | OTUs in common | OTUs unique: ethanol | OTUs unique: bulk | Total OTUs |
| --- | --- | --- | --- | --- | --- |
| Complete dataset | 6 | 357 | 640 | 104 | 1101 |
| Complete dataset * | 6 | 301 | 115 | 98 | 514 |
| Macroinvertebrates* | 6 | 262 | 39 | 97 | 398 |
| Macroinvertebrates subsample* | 4 (excl. S2/S4) | 232 | 39 | 85 | 356 |
| EPTD taxa* | 4 (excl. S2/S4) | 162 | 16 | 35 | 213 |
| EPT taxa* | 4 (excl. S2/S4) | 46 | 4 | 10 | 60 |

\* OTUs > 85 % similarity to a BOLD entry only

**Figure 3:**
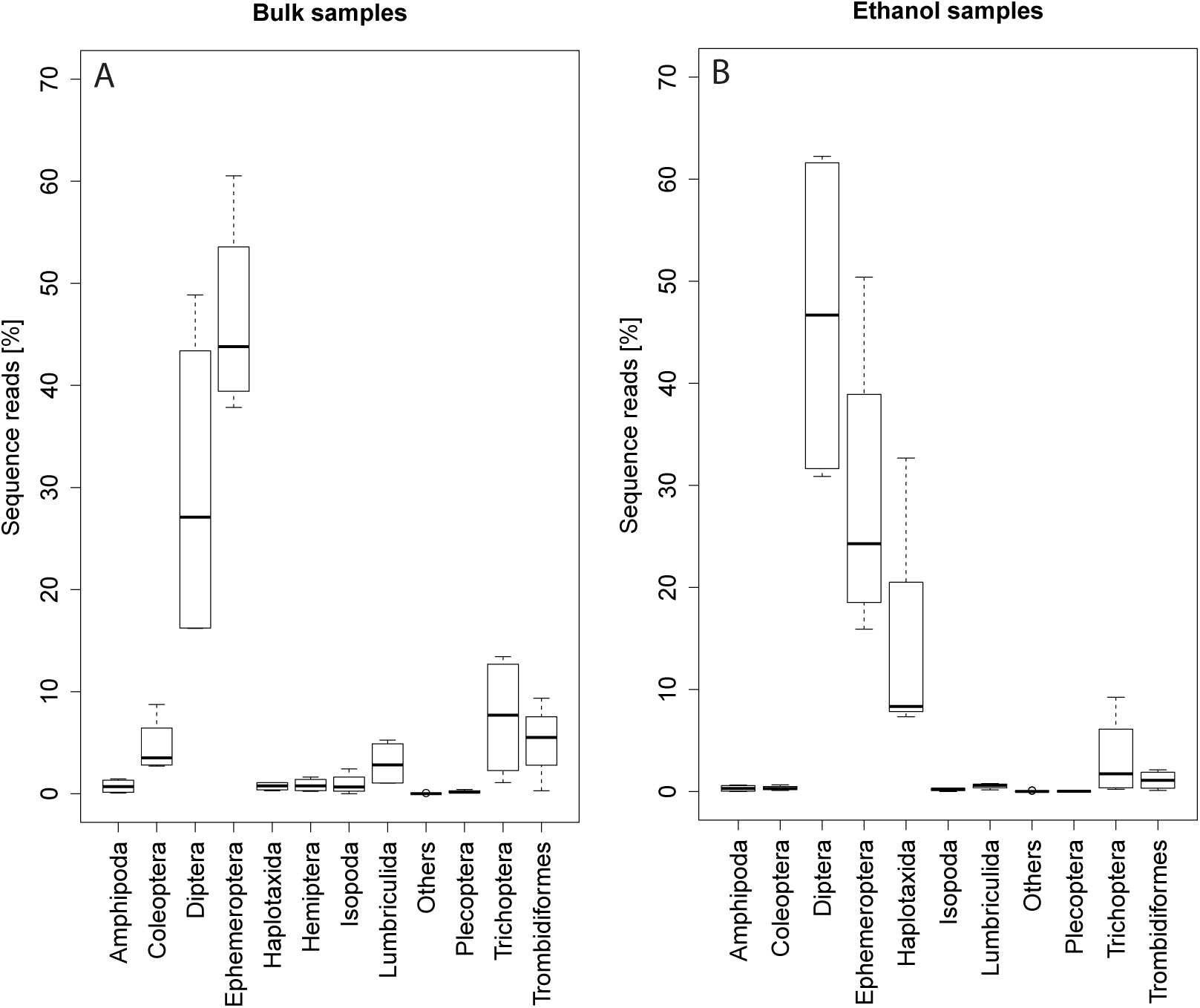
Proportion of reads per order from all macroinvertebrate taxa in A) bulk samples and B) ethanol samples. The category “Others” includes all orders with read numbers < 0.5 %.

Including only OTUs assigned to Arthropoda, Annelida, Mollusca or Platyhelminthes with at least 85 % similarity to records found in BOLD, ethanol samples included on average 472,688 (± 494,305, S1E-S6E) reads, while bulk samples contained on average 384,261 (± 74,398, S1B-S6B) reads (Fig. 2, Tab. S9). Again comparing this filtered data set, 262 OTUs were shared between the two methods, while 39 were unique to the ethanol and 97 to the bulk samples, respectively. Out of the 42 OTUs assigned to beetle taxa in bulk samples, only 11 were detected in ethanol samples, which showed in addition extremely low read numbers (Fig. 3, Tab. S9). On the other hand, 12 OTUs belonging to the Clitellata (mainly Haplotaxida) were only found in ethanol samples (Tab. S9). Due to high differences in read numbers per sample (Fig. 2), a sample subsetting was performed retaining only 230,000 reads identified as macroinvertebrates per sample (Tab. S10). The samples S2E and S4E showed macroinvertebrate read number < 50,000 and where therefore excluded from further analysis. For comparisons to bulk samples, also S2B and S4B were excluded. Comparing macroinvertebrate data between the two methods based on the four remaining samples, both datasets had 232 OTUs in common, 39 were unique to the ethanol and 85 to bulk samples, respectively (Tab. 2).

Reducing the taxonomic focus further (subsampled dataset) to Ephemeroptera (mayflies), Plecoptera (stoneflies) and Trichoptera (caddisflies), the overlap of both methods was 46 OTUs, four OTUs were unique to the ethanol samples (Ephemeroptera: *Ephemerella* sp. Trichoptera: *Goera pilosa*, *Hydropsyche contubernali*, *Anabolia nervosa*) and 10 to the bulk samples (Ephemeroptera: *Habrophlebia lauta*, *Ecdyonurus torrentis*, *Baetis muticus*, *Leuctra albida*; Trichoptera: *Oecetis testacea*, *Sericostoma flavicorne* (2x), *Limnephilus lunatus*) (Tab. S11). Average read numbers were higher in bulk samples for all three EPT taxa (Ephemeroptera ethanol: 66,055, bulk: 106,924; Plecoptera: 68.5, 448.5; Trichoptera: 8487.5, 17190). Including taxa belonging to the Diptera (EPTD) in both methods 162 identical OTUs were detected. Thereby 16 OTUs were unique to the ethanol tests and 35 to the bulk samples. Within those OTUs, 12 uniquely found in bulk samples belonged to the Chironomidae, while eight OTUs assigned to chironomids were only found in ethanol tests (Tab. 2).

Including either all detected OTUs after the quality filtering steps (1101 OTUs) or those with > 85 % sequence similarity to a BOLD record (514 OTUs), PCoA analyses (Fig. 4A and B) using similarity measures on abundance (Bray-Curtis) and presence/absence (Sørensen) data show clearly distinct community compositions for all sample sites and especially between the two sources of DNA (ethanol vs. bulk). While the variance associated to the first axis is clearly and almost exclusively related to the source of DNA based on presence/absence data (Fig. 4 A and C), the same signal is recovered by axis one and two when read abundance numbers are included (Fig. 4 B and D).

**Figure 4:**
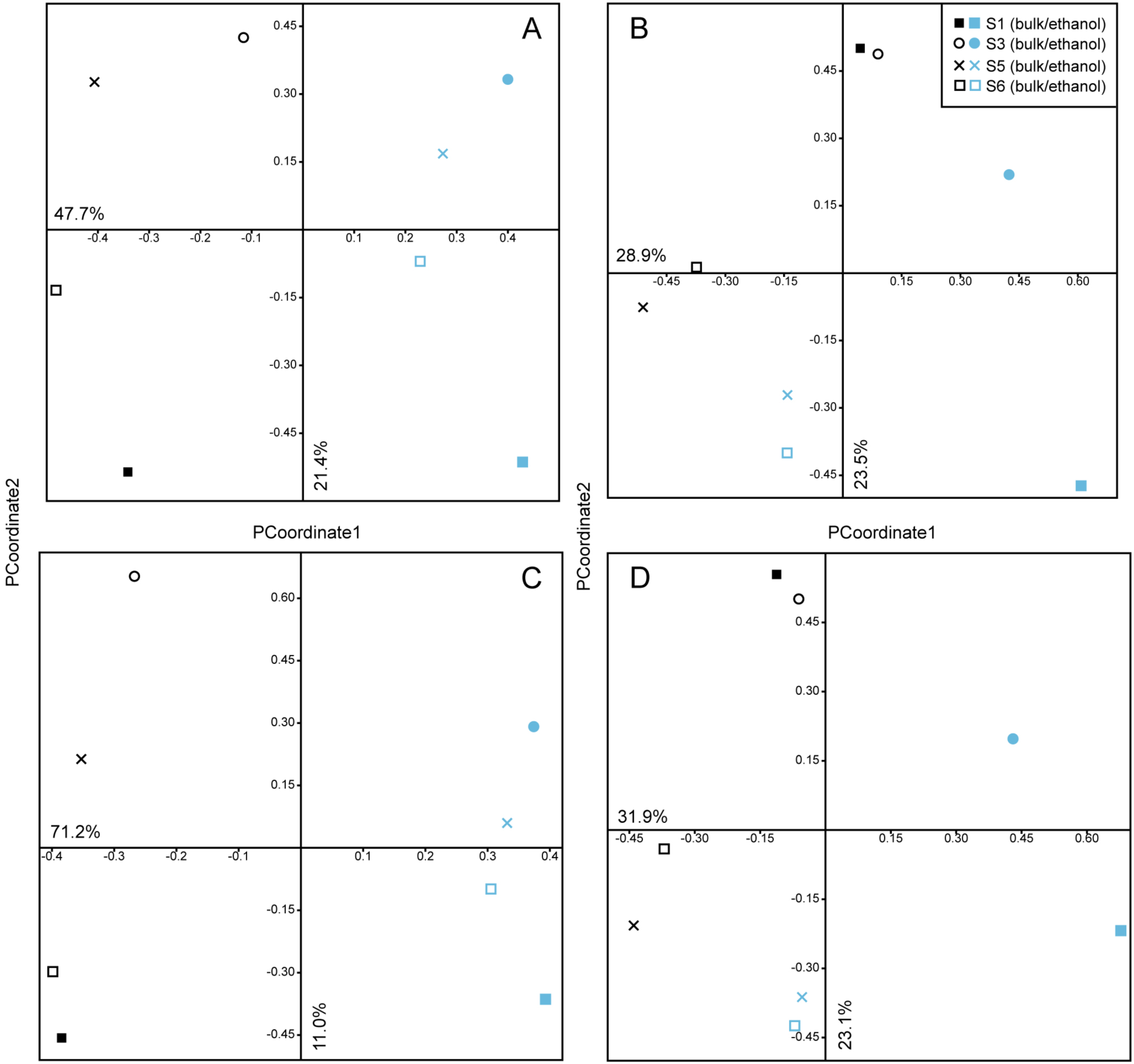
Visualization of differences in community composition with scatter plots from PCoA for the four retained sample locations S1, S3, S5, S6 obtained via bulk and fixative metabarcoding using the ‘reduced’ data set (A & B, 514 OTUs above 85% similarity database hits) and the complete data set (C & D, 1101 OTUs, no similarity threshold filtering). A and C show data based on presence-absence similarity, B and D include read abundance information.

When comparing the composition of OTUs belonging to macroinvertebrate EPTD taxa between the sample sites, the effect of the two different sources of DNA becomes smaller and the sites cluster closer together in the scatter plots (Fig. 5) than compared to the results from the full OTU comparisons (Fig. 4). Using abundance information (Bray-Curtis similarity, Fig. 5 B), the communities from each sample point appear to become less similar in the scatter plot, with S1 being most strongly affected.

**Figure 5:**
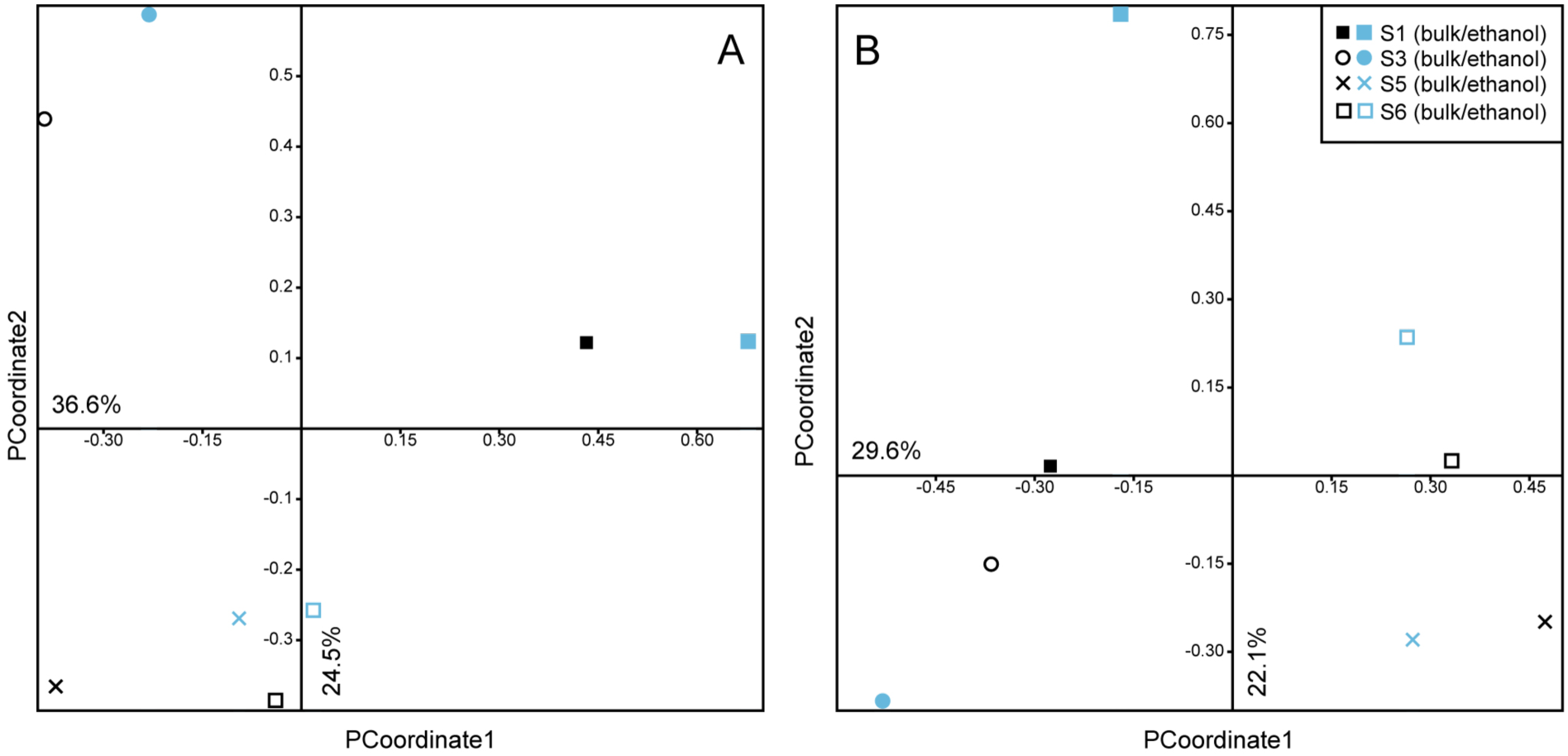
Visualization of differences in EPTD taxa community composition with scatter plots from PCoA for the four retained sample locations S1, S3, S5, S6 on the 213 OTUs (taxa with >85% similarity to BOLD database entries) and based on presence/absence (A) and read abundance (B) data.

## 4. Discussion

### 4.1 Testing filtration of ethanol and isolation of DNA from filters

The extraction of DNA from the fixative worked for three (protocol 0, 2 and 3) of the four tested protocols and an analysis of sequencing results was possible. The insufficient sequencing output of samples processed with ultrasonic irradiation (protocol 1) could be due to the low irradiation frequency that was used. The applied ultrasonic water bath (Bandalin SONOREX Super RK 510 H) works at a predetermined frequency of 35 kHz with indirect irradiation only. We chose this approach in order to exclude potential cross contamination through an ultrasonic probe system with direct contact between the device and sample material. If direct ultrasound irradiation at higher frequencies would induce an increase in cell disruption and thus higher rates of release of DNA to the fixative (e. g., Sinisterra 1992, Rokhina 2009, Kwiatkowska et al. 2011) should be tested in subsequent studies.

Taxa detection in samples processed with protocol 0 (first ethanol phase, no treatment), protocol 2 (shaking) and protocol 3 (freezing) were highly similar, with mollusc taxa most consistently undetected. Especially the snail *Ancylus fluviatilis* was rarely detected and if, contributed only low read numbers. This freshwater limpet is of small body size (in the present study on average 4 mm) and dorsally covered by a shell and only a small body fraction is free from protection, which can in addition be covered by a weak layer of mucus (Calow 1974). Similar characteristics apply to the other mollusc taxa in the mock communities (identified only to Gastropoda in the field). As revealed by standard DNA barcoding and reverse identification via BOLD the representatives in our samples belonged to seven different taxa. The four species *Potamopyrgus antipodarum*, *Physa acuta, Radix labiata* and *Succinea putris* where successfully detected using metabarcoding based on moderate read numbers (average 0.2 %). The other taxa (*Galba truncatula*, *Physa* sp. and Succineidae) were rarely detected and no similarity between sequences generated by single-specimen barcoding and sequences in the OTU table was given. However, even if the comparison between single-specimen barcoding and metabarcoding was problematic, results indicate that the detection of molluscs through fixative-based metabarcoding is not reliable. In comparison, detection failures for arthropod species in our mock communities were not consistent. This was most noticeable for the samples processed with protocol 0, where certain taxa were randomly not detected (community A2: *Habroleptoides confusa*, C2: *Polycentropus flavomaculatus* and *Hydropsyche saxonica*, C3: *Ephemera danica*, E1: *Ecdyonurus torrentis*). For protocol 2 (shaking) additionally to *Ancylus fluviatilis*, *Polycentropus flavomaculatus* could not be detected in three different communities, while protocol 3 (freezing) left only the two mollusc taxa *Ancylus fluviatilis* and Succineidae sp. undetected. Additionally, the number of reads assigned to the eight taxa of interest was the highest and most consistent (93.7 % ± 3.5) for protocol 3 (freezing), which can be considered as the most successful protocol to increase the DNA release of specimens to the fixative.

In addition to the eight taxa in the mock communities, a high proportion (on average 19.12 %) of the total reads was assigned to other macroinvertebrates in samples treated with protocol 0 (9.35 % and 2.42 % in protocols 2 and 3, respectively). The resulting OTUs matched various dipterans, but also Clitellata and Ephemeroptera and a few other arthropod taxa (Tab. S6). In addition to particles or cells attached to the mock community specimens, stomach content of predatory taxa (e. g., *Gammarus*, *Hydropsyche*, *Polycentropus*) might be the source of this additional diversity. This assumption is supported by the fact that most of the additional OTUs belong to taxa, which frequently serve as prey (mainly dipterans) (Cummins 1973, Klecka and Boukal 2013). Furthermore, it can be observed that some specimens regurgitate their stomach content when being preserved, which consequently should be detected by fixative-based metabarcoding. For protocol 2 and 3 a lower proportion of reads was assigned to these OTUs with potential stomach content origin, most likely because specimens were already dead when transferred to the second ethanol phase and did not release stomach content into the fixative.

Only for protocol 0 a weak correlation between biomass and read numbers existed (Spearman’s rho 0.211, *p*=0.02). The strong scattering of the underlying data points and the fact that no significant correlation between biomass and read numbers was evident in the two other treatments indicate that biomass of individuals has an inferior effect on the final read number. However, the present results show that individuals release different amounts of DNA, which biases in turn the generated read numbers. For fixative-based metabarcoding, these biases seem to be more dependent on morphology of the specimens or on the random dispersal of cells or even body structures on the filter. An additional filtering with a finer mesh size than applied in the present study (0.5 mm) of the ethanol before processing could help circumventing the problem of complete cells on the filter, as recently also proposed by a comparison of different filtration techniques for eDNA from water samples (Majaneva et al. 2018).

To accomplish higher DNA yield and thereby increase taxa detection including the stomach contents of specimens for a more comprehensive community assessment, the combination of protocol 0 and 3 might be optimal. This combination does also decrease workload, because it excludes the ethanol change step and only includes the transfer of samples to liquid nitrogen or an ultra-low temperature freezer (- 150°C) after sampling for a determined time (e. g., one hour).

### 4.2 Comparison of environmental bulk samples with ethanol samples

A comparison between bulk and ethanol samples based on all detected taxa revealed completely different communities (Fig. 4). In total, 640 OTUs were unique for the ethanol derived samples, which were mainly composed of OTUs assigned to algae (Bacillariophyta, Ochrophyta, Rhodophyta) and bacteria (Proteobacteria), but also Amoebozoa and Porifera (Tab. S8). These groups are mainly present in ethanol samples, most likely because whole kick net bulk samples include algae, parts of wood, gravel and other substrate transferred to the ethanol and consequently released DNA into the fixative, or function as substrate for other groups (bacteria etc.). The degeneracy of the used BF2/BR2 primers leads to the amplification of even not closely related taxa other than macroinvertebrates and might explain the high abundance of “untargeted taxa” in the dataset (Elbrecht et al. 2017, Macher and Leese in review). This phenomenon is also known from eDNA (environmental DNA) studies where, next to the taxa of interest a lot of other groups were present in final sequencing results (Deiner et al. 2015, Macher and Leese in review). While for the sample points S3E, S5E and S6E the riverbed substrate was dominated by gravel and stones, for sample points S1E and S2E substrate was overgrown with algae. For these two sample points and especially S2E, read numbers were dominated by algal OTUs and only a minor fraction of reads was assigned to macroinvertebrate taxa (Fig. 2). Sample S2E was even excluded from further analysis due to the very low read numbers. In such situations, a brief separation of organisms from larger plant material or a preceding filtering step would presumably decrease the number of reads assigned to “untargeted taxa”. Furthermore, if a particular taxonomic group is of interest, the usage of more specific primers would increase amplification accuracy (Elbrecht and Leese 2017). For the samples S4E and S6E problems during PCR were probably due to an increase in inhibiting substances at these sample sites. In comparison to ethanol samples, extraction of DNA from the bulk samples was conducted with macroinvertebrate organisms only, which were separated from any substrate and additionally sorted in two size categories. As illustrated in Figure 2, the distribution of total read numbers among samples is more consistent and the majority of reads is assigned to macroinvertebrate taxa.

Due to the large differences in read numbers assigned to macroinvertebrate taxa within ethanol samples (Fig. 2), but also in comparison to bulk samples, a rarefaction of 230,000 reads was performed. For rarefied samples 232 OTUs were shared between ethanol and bulk samples, while 39 and 85, respectively, were unique to one of the methods. Regarding read numbers, the orders Diptera (mainly chironomids) and Haplotaxida were stronger represented in ethanol samples (Fig. 3) and for the latter also OTU richness was higher in ethanol (36) than in bulk samples (26). Both orders comprise taxa with soft body structures and surface, where only the head is partly sclerotized. This can entail an increased DNA release of these taxa compared to other organisms with a more pronounced skeleton. On the other hand, many chironomids as well as oligochaetes serve as prey for other macroinvertebrates (Hildrew and Townsend 1982, Krisp and Meier 2005, Klecka and Boukal 2013) and the additional OTUs or reads could result from DNA of regurgitated prey (see also above). The detection of prey organisms would be a positive by-product of this protocol, since it provides a more detailed insight into the present community. For bulk sample metabarcoding approaches, we speculate that the detection of prey organisms remains difficult unless a much higher sequencing depth is applied, due to the lower proportion of prey DNA in stomachs as compared to the actual specimen tissue.

The high number of detected chironomid OTUs (>80) and assigned species names to them (62 with >98% sequence similarity) in the ethanol derived DNA is especially promising, as these non-biting midges are ecologically highly diverse and dominant in many aquatic ecosystems (Ferrington 2008). Chironomids have therefore been proposed as indicator group for freshwater ecosystem quality, but as their morphological identification is extremely difficult they are not used for Water Framework Directive monitoring in Germany (cf. Operationelle Taxaliste). These difficulties and the lack of taxonomic experts for chironomids are also the reason for the low coverage of species with DNA barcodes available (ca. 25 % of 700 species in Germany; www.bolgermany.de). Our non-destructive approach for metabarcoding benthic communities might constitute a promising additional source for reference chironomid species, which can either be studied morphologically after having been detected and taxonomically assigned, or which can indicate specific sample locations for emergence trapping (as usually the adult males are needed for an unambiguous morphological identification). The potential of this approach is illustrated by the fact that 29 of the 62 via BOLD to species level identified chironomids are still missing in the German Barcode of Life reference database (as of March 21^st^ 2018) and where thus identified by DNA sequence comparisons to material from other countries. Theissinger et al. (2018) recently showed how valuable species level knowledge on chironomids would be, as several species are indicators for good water quality (low saprobic values), but are most often lumped together as *Chironomus* sp. with a high saprobic value indicative for bad water quality.

As mentioned above, bulk sample metabarcoding revealed a higher OTU diversity assigned to macroinvertebrates (317) and also read numbers for Amphipoda, Coleoptera, Isopoda, Lumbriculida, Trichoptera and Trombidiformes were higher than in the ethanol samples (Fig. 3). We speculate that this is due to a lower relative number of template DNA fragments for the PCR for the aforementioned taxa in the DNA mixture from the filters. Representatives of these taxa often have a pronounced exoskeleton or are surrounded by a case of stable material (trichopterans), which can lead to the restraint of DNA. Additionally, Trombidiformes are extremely small, which further reduces DNA release. The decreased number of reads available for those taxa might further be explained by the competition for sequencing coverage, especially when high fractions are “used up” for other taxonomic groups (algae etc., see above). However, regarding only EPT taxa (Ephemeroptera, Plecoptera, Trichoptera) the two methods provided 46 overlapping OTUs with an additional 4 OTUs unique to the ethanol samples and 10 OTUs unique to bulk samples. Within the Water Framework Directive 2000/60/EC these taxonomic groups are essential biological quality elements (BQEs) for the assessment of the ecological status of rivers and lakes (Meier et al. 2006). With 50 OTUs (40 different species determined) detected through the extraction of DNA from the fixative and subsequent metabarcoding and only 6 OTUs less than with metabarcoding bulk samples. These results indicate the utility of this method for biodiversity assessment of certain taxa or for biomonitoring approaches without the time-consuming step of specimen sorting and the consequent decrease in costs compared to bulk sample metabarcoding or morphological identifications. Especially in cases where legal frameworks do not allow homogenizing the organismal samples, metabarcoding of the fixative could become a viable alternative.

## 5. Conclusions

We showed that metabarcoding based on DNA extraction from fixative (ethanol) is in general possible, although several false negative cases are to be expected (especially mollusc taxa). The controlled mock-community, as well as the field sample analyses showed that detection of biological quality elements relevant for ecological status class assessment (EPTD) was highly successful and results of fixative metabarcoding was comparable to taxa lists generated by bulk sample metabarcoding. The extraction of DNA from ethanol and the subsequent metabarcoding is an interesting less-invasive and time efficient alternative to standard metabarcoding approaches and at the same time shows biodiversity detection comparable to morphological approaches.

## Acknowledgement

VZ, FL, BP and MFG are members of the German Barcode of Life (GBOL) project, which is generously supported by a grant from the German Federal Ministry of Education and Research (FKZ 01LI1101 and 01LI1501). We want to thank Jan-Niklas Macher and Vasco Elbrecht for support in study design and helpful discussions, Laura von der Mark and Jana Thormann for their help in the laboratory and the Leese lab journal club for proofreading this manuscript.

## Authors contributions

FL, MFG and VZ conceived the ideas and designed the methodology; VZ and BP carried out the laboratory work and VZ performed bioinformatic analyses; VZ led the writing of the manuscript. All authors contributed critically to the drafts and gave final approval for publication.

## Supplements

**Script S1:** R script for NGS data analysis as implemented in JAMP

**Table S1:** Assembly of 15 mock communities with size relations and treatments listed.

**Table S2:** Read numbers for different mock community samples after demultiplexing.

**Table S3:** Comparisons of sequences generated through single-specimen barcoding with the database BOLD, taxonomic assignment (species, similarity, database ID) and comparison with metabarcoding results. Proportion of reads for each taxon in OTU table as well as the OTU number is given.

**Table S4:** Biomass (g) and length (mm) for each specimen in mock communities.

**Table S5:** Complete OTU table of mock community analysis.

**Table S6:** OTU table of mock community analysis with the eight targeted taxa excluded.

**Table S7:** Read numbers of ethanol and bulk samples after demultiplexing.

**Table S8:** OTU table of ethanol and bulk sample analysis.

**Table S9:** OTU table of ethanol and bulk sample analysis with only macroinvertebrates included.

**Table S10:** OTU table of ethanol and bulk sample analysis with only macroinvertebrates included and a subsampling of 230,000 reads.

**Table S11:** OTU table of ethanol and bulk sample analysis with only EPTD taxa (Ephemeroptera, Plecoptera, Trichoptera, Diptera) included.

